# Identifying interaction interface between *Pseudomonas* major biofilm forming functional amyloid FapC and disordered chaperone FapA during fibrillation deceleration

**DOI:** 10.1101/2023.03.14.530334

**Authors:** Chang-Hyeock Byeon, Hakan Saricayir, Kasper Holst Hansen, Jasper Jeffrey, Maria Andreasen, Ümit Akbey

## Abstract

Functional bacterial amyloids (FuBA) play a crucial role in the formation of biofilms, which are mediating chronic infections and contribute to antimicrobial resistance. This study focuses on the FapC protein from *Pseudomonas*, a major contributor to biofilm formation. We investigate the initial steps of FapC amyloid formation and the impact of the chaperone-like protein FapA on this process. Using solution NMR spectroscopy, we show that both FapC and FapA, which are part of the same biofilm-forming protein operon, are intrinsically disordered proteins (IDPs) in their soluble monomeric state. These SSPs were determined and compared to the Alphafold models. We further demonstrate that the IDP chaperone FapA interacts with FapC and significantly slows down the formation of FapC fibrils, while maintaining the fibril morphology unchanged. Our NMR titration experiments reveal that ∼18% of the resonances show FapA induced chemical shift perturbations (CPSs) which has not been previously observed, the largest being for A82, N201, C237, C240, A241 and G245 residues. These sites may suggest a specific interaction site and/or hotspots of fibrillation inhibition/control interface at the R1/L2 and L2/R3 transition areas and at the C-terminus of FapC. Remarkably, ∼90% of FapA NMR signals exhibit substantial CSPs upon titration with FapC. A temperature dependent effect of FapA was observed on FapC by ThT and NMR experiments. This study provides a detailed understanding of the interaction between the chaperone/chaperone-like FapA and the functional amyloid protein FapC, shedding light on the regulation and slowing down of amyloid formation. Our findings have important implications for the development of therapeutic strategies targeting biofilms and associated infections, leveraging these structural and mechanistic insights.

## Introduction

Functional bacterial amyloids (FuBAs) constitute a unique class of amyloid fibrils that possess functional properties in living organisms, distinct from their pathological counterparts that cause diseases.^1-6^ Their involvement in various biological processes, such as biofilm formation, enables bacteria to survive and protect themselves under stressful conditions. While FuBAs themselves do not directly induce diseases, they indirectly contribute to pathology through chronic bacterial infections, thereby leading to increased antimicrobial resistance. Chronic infections, often caused by biofilm-forming bacteria, account for a significant portion of such cases.^7^ Several biofilm-forming FuBAs, including CsgA from *E. coli*, FapC from *Pseudomonas*, TasA from *B. subtilis*, and PSMs from *S. aureus*, have been extensively studied,^8-11^ summaries in recent reviews provide detailed insights.^6, 12^

Recent years have witnessed a growing interest in studying functional amyloids, elucidating their biological roles, and characterizing their biophysical properties. Despite the abundance of biophysical information available, the structural understanding of amyloid formation, from monomeric IDP to fibrillar filamentous forms within biofilms, remains elusive.^13^ This lack of structural information hinders the development of therapeutic strategies against biofilm-mediated chronic infections and antimicrobial resistance. To overcome this obstacle in the fight against biofilm-mediated chronic infections and/or biofilm-mediated antimicrobial resistance, structural insights obtained through novel approaches are essential.

Many functional and pathological amyloid-forming proteins exist in monomeric forms as intrinsically disordered proteins (IDPs), such as CsgA, FapC, αSynuclein, and amyloid β, while examples of folded proteins, including the functional amyloid TasA and pathological amyloids insulin and transthyretin, also exist.^6, 14^ Studying monomeric prefibrillar protein forms, particularly the IDP ones, poses challenges and requires techniques capable of achieving atomic resolution.^15-17^ High-resolution solution nuclear magnetic resonance (NMR) spectroscopy emerges as a powerful option, offering atomic-level insights into the first steps of oligomerization, soluble monomers, soluble oligomers, and other high-molecular-weight soluble proteins.^18, 19^ In contrast, commonly used methods like Thioflavin T (ThT) assays detect bulk fibrillation but lack atomistic information.^20^ Solution NMR has successfully characterized NMR-visible soluble monomeric species, NMR-invisible high-molecular-weight oligomeric/protofilament species, and sparsely populated oligomeric species of various amyloid fibrils, both functional and pathological.^18, 21-25, 26-30^

To address the structural knowledge gap concerning functional amyloids, this study aims to provide insights into the molecular structure and organization of functional amyloids. We focus on the prefibrillar soluble monomeric form of the functional amyloid-forming protein FapC from *Pseudomonas* using solution NMR spectroscopy.^31^ Fibril formation strongly depends on factors such as protein concentration, temperature, and sample nature (monomeric state versus a mixture of monomer/oligomer). While higher protein concentrations benefit NMR measurements by enhancing overall signal observation, they also accelerate fibrilization rates, making timely observation challenging. Temperature significantly influences fibrillation rates, and the presence of high-molecular-weight species, such as seeds or soluble/insoluble oligomers at the start of fibrillation, affects the process and may introduce biases if not addressed properly.^32^ These factors were carefully considered in this study.

Chaperones have the ability to modulate the fibrillation of amyloid fibrils, exhibiting varied effects ranging from slowing down fibril formation to prevention or even reversal.^17, 21, 23, 33-35^ Examples from both pathological and functional amyloid perspectives demonstrate a wide range of chaperone effects on fibrilization. For instance, CsgC slows down fibril formation of CsgA functional amyloid,^23^ while HSP90 family chaperones delay the pathological amyloid fibrillation of αSynuclein.^33^ FapA, an intrinsically unfolded protein and a chaperone from *Pseudomonas*, belongs to a unique class of proteins that can act as chaperone or chaperone-like.^17^ Understanding the molecular mechanism of FapA chaperone can aid in developing strategies to prevent amyloid formation in the context of biofilms and related chronic infections. In this work, we present a time-resolved, high-resolution solution NMR approach to characterize the prefibrillar monomeric form of FapC and its transition toward amyloid fibril formation. As an initial step toward achieving a complete molecular understanding of biofilm-forming FuBAs, we monitored changes in the NMR spectrum of monomeric FapC. By utilizing our recent NMR resonance assignment,^31^ we determined time-dependent chemical shift and intensity changes, which were correlated with FapC secondary structure to comprehend the first interaction interface. Additionally, we quantified the impact of the IDP chaperone FapA on FapC fibrillation, aiming to establish a general chaperone-based mechanism for amyloid fibril prevention/slowing down. To prevent seeding/oligomerization effects, low protein concentrations and freshly prepared FapC samples were used. Furthermore, NMR experiments were conducted at a lower temperature of 274 K to decelerate fibrillation kinetics, avoiding transient effects associated with fibrillation and enabling observation of all FapC NMR resonances, which partially disappeared at elevated temperatures under the current experimental conditions.

## Results

### FapC is an intrinsically disordered protein

By utilizing solution NMR spectroscopy, we recently presented the complete NMR resonance assignment of soluble monomeric FapC (BMRB #51792-51793).^31^ NMR spectra and secondary structure propensities (SSPs) determined using NMR chemical shifts indicate that FapC is an intrinsically disordered protein (IDP) with still significant α-helix and β-sheet structural propensities,^6^ see **Figure 1A,E**. The 2D ^1^H-^15^N HSQC NMR spectra shown in **Figure 1 A**,**B** of both the pure FapC and the FapC complexed with its chaperone FapA at 1:4 mole ratio show a narrow proton chemical shift dispersion of ∼1 ppm, due to their IDP nature. Detailed sequence properties of FapC and FapA based on FELLS analysis are shown in **Figure** 1 **– figure supplement 1**.^36, 37^ **Figure 1E** depicts the overlay of FapC domain organization of FapC (L: loop, R: repeat, C: c-terminus and N: n-terminus), the NMR determined SSPs and the blown-up representation of the β-aggregation potential (aggregation prone regions, APRs) predicted using three algorithms TANGO, WALTZ and AGGRESCAN.^38-40^ The normalized plots of these predicted APRs are shown in **Figure 1 – figure supplement 2**. Compared to the domain organization consensus, these *in silico* methods indicate slightly different APRs, however, the regions between amino acids 39-45 (∼5% propensity by TANGO algorithm), 102-109 (∼46%), 142-148 (∼1.2%), 156-163 (∼18%), 197-205 (∼0.1%), 215-222 (∼1%) are covered by all three algorithms though with varying propensities. These APRs correlate with the three repeat regions of FapC (R1,R2 and R3), but also falls in the unexpected areas outside the repeats in the N, L1 and L2 of FapC. This observation of APRs in the non-repeating FapC regions fits to our previous observation that L1/L2 loop regions increase amyloid formation rates determined by ThT assays.^37^ The NMR determined SSPs of the FapC indicates predominantly random coil with up to ∼40% polyproline 2 (PP2) helix conformations. This large PP2 propensity could be important for structural transition towards the correct amyloid fibril fold in contrast to amorphous aggregates.^41^ Nevertheless, sizeable β-sheet propensities between 5-20% are observed throughout the protein, along with a few α-helix indicating areas 181-189 with ∼15% and 113-121 with ∼7% propensity, based on the ncSSP analysis shown in **Figure 1D**. There are areas that indicate turns or helices with smaller propensities, such as residues at around 37, 57-60, 70, 87-90, 151-155, 228, 236 and 245-247. The predicted APRs fall within the NMR determined β-sheet propensity areas, indicating a good correlation between the NMR and *in silico* methods. The structural transition from the monomeric to amyloid FapC during primary nucleation could be initiated at/around these APRs. Overall, even though the NMR indicate relatively small SSPs, they correlate well with the secondary structure elements predicted by the Alphafold model, shown in **Figure 2A**. This highlighting a possible less hindered conformational transition from monomeric IDP fold into the amyloid FapC fold.

**Figure 1.**
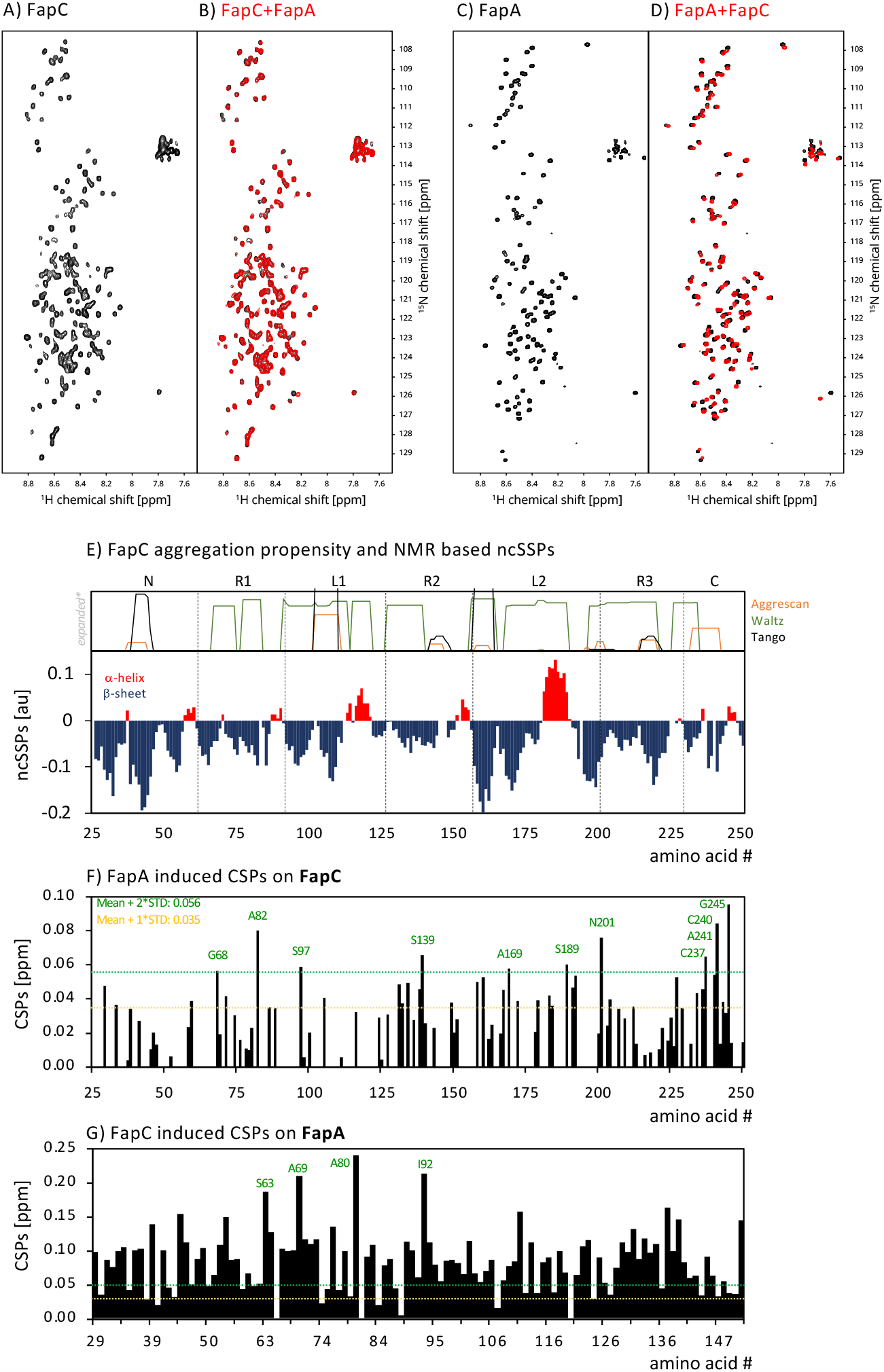
**A)** 2D fingerprint ^1^H-^15^N HSQC NMR spectra of pure full-length ^15^N-labelled FapC (black), **B)** ^15^N-labelled FapC mixed with unlabeled FapA (red) at 1:4 molar ratio (FapC+FapA), overlayed with spectrum in black from A. **C)** 2D fingerprint ^1^H-^15^N HSQC NMR spectra of pure full-length ^15^N-labelled FapA (black), **D)** ^15^N-labelled FapA mixed with unlabeled FapC (red) at 1:4 molar ratio (FapA+FapC), overlayed with spectrum in black from C. The spectra are shown at the identical contour levels for direct comparison. The NMR resonance assignment for FapC is presented previously.^31^ FapC and FapA concentration for pure samples is 50 μM, and the corresponding FapA and FapC concentrations are adjusted accordingly for each sample at a 1:4 molar ratios. The 2D spectra were all recorded at 274 K. **E)** Representation of amyloid forming regions predicted by three different algorithms, along with the NMR chemical shift-based neighbor-corrected secondary structure propensities (ncSSPs) for the full-length FapC. Y-axis value of 1 refers to 100% propensity for ncSSPs. Note that the scale for amyloid prediction plot is not up to 100%, and the amyloid regions are expanded for clear visibility and to highlight even a small amyloidogenic region. The correctly normalized amyloid propensities are shown in Figure 1 – supplement 2. The ncSSP scale is normalized, and 1 corresponds to 100% propensity. **F)** Representation of the FapA induced chemical shift perturbations (CSPs) in the FapC NMR spectrum at the FapC:FapA 1:4 sample compared to the pure FapC at one day of incubation. The description of the significance in the results parts. The labels are shown only for the CSPs larger than “mean + 2*STD”. **G)** Representation of the FapC induced chemical shift perturbations (CSPs) in the FapA NMR spectrum at the FapA:FapC 1:4 sample compared to the pure FapA at one day of incubation. Note that the CSPs in the FapA spectrum is much larger compared to the CSPs in FapC spectrum. The weighted cumulative CSPs given in terms of ppm (y-axis) and are calculated based on Δ δ = [(5*(δ ^1H^_FapC:FapA_ – δ ^1H^_FapC_))^2^+ (δ^15N^_FapC:FapA_ – δ^15N^_FapC_)^2^]^0.5^ and are 0.015 and 0.087 ppm for FapC:FapA and FapA:FapC samples, respectively. **Figure supplement 1**. FELLS analysis of FapC and FapA **Figure supplement 2**. Normalized plots of TANGO, WALTZ and AGGRESCAN aggregation potentials for the aggregation prone regions (APRs).

**Figure 2.**
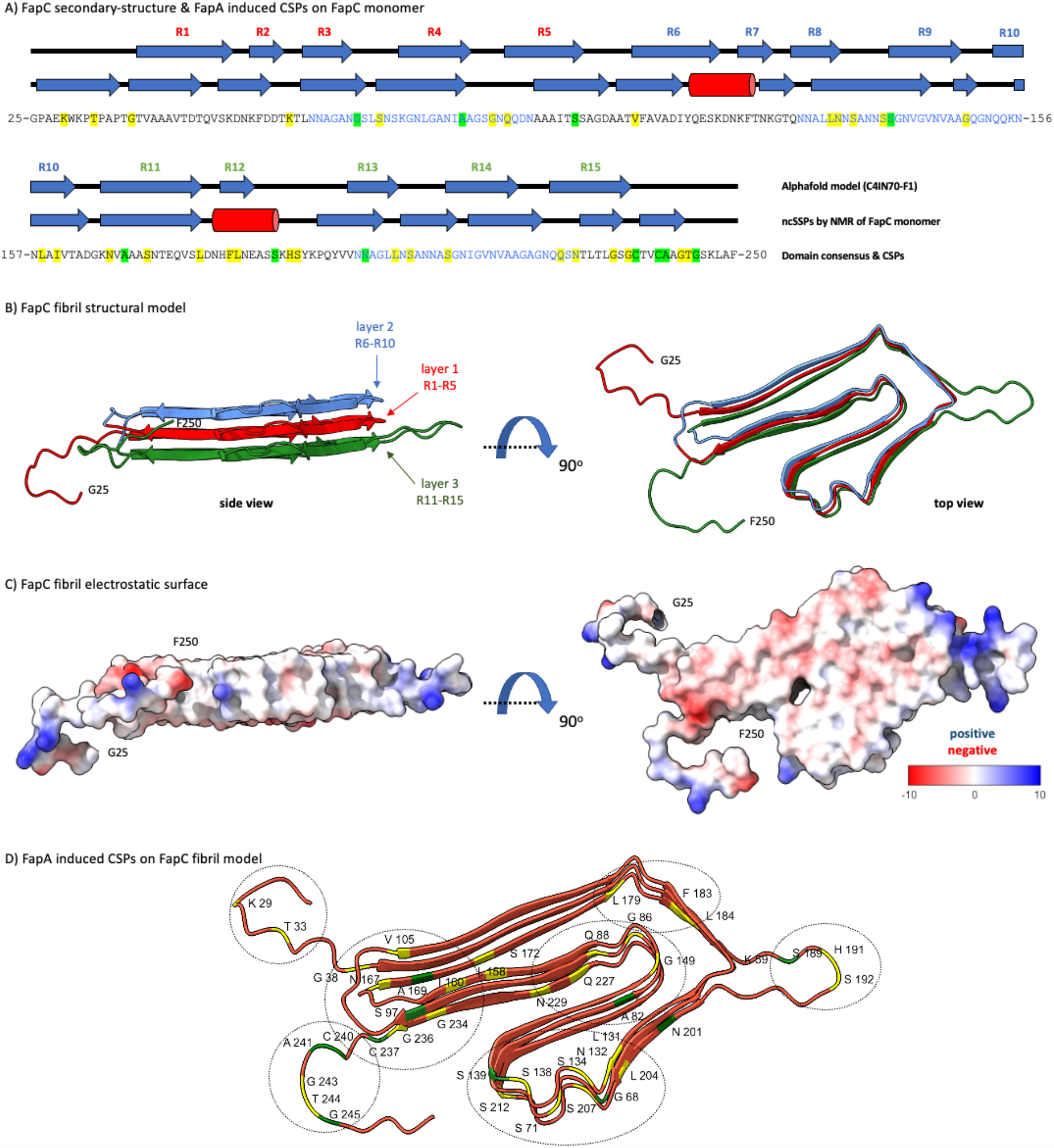
**A)** The secondary structure comparison of the amyloid FapC structure based on Alphafold model (AF-C4IN70) and the neighbor-corrected secondary structure propensities (ncSSPs) based on our experimental solution NMR chemical shifts of the monomeric FapC with the BMRB entry #51793. The amino acid sequence of FapC is shown at the bottom. The black color represents the N-terminus, loop regions and C-terminus in the sequence, whereas the blue color represents the generally accepted repeat regions (R1-R3). The Alphafold secondary structure elements are depicted by the model. We used a threshold of 4-residues for depicting the α-helices (red cylinders) or β-sheets (numbered and blue arrows) as a secondary structure element for the NMR ncSSPs data. The yellow (significant CSPs) and green (more significant CSPs) highlights are following the notation shown and described in Figure 1F. The FapC sequence (between 25-250 amino acids) is shown to visualize the CSPs, the repeat regions are colored in blue. Yellow highlight indicates significant changes, and the green highlight indicates much larger changes as depicted with two dashed lines in the CSPs plot. **B)** Representation of Alphafold amyloid FapC structural model in two orientations, side, and top view (note that this is a model for the amyloid fibril structure of FapC). The residues 1-24 are omitted in the model, following the FapC construct studied here. The repeats forming three different layers are indicated with color coding: beta-strands R1-5 form layer 1 (red), beta-strands R6-10 form layer 2 (blue) and the beta-strands R11-15 form the layer 3 (green). Note than the nomenclature used for the FapC protein segments (repeat region 1-3) are different from the beta-strand numbering here. **C)** Representation of the electrostatic surface for the FapC fibrillar structure model shown in B. The electrostatic surface is produced by automatically determined coulombic values in terms of kcal/(mol*e) and with minimum:-17.78, mean:-0.75, and maximum:13.51 values, by using Chimera X. **D)** Locations of the chemical shift perturbations (CSPs) on the structural model, that are observed in the solution NMR spectra of isotope labeled monomeric FapC as a soluble IDP upon addition of four-fold unlabeled FapA chaperone. The green and yellow colors are placed according to the representation in Figure 1E, represent the large (green) and medium (yellow) CSPs. The chemical shift perturbation should be understood with care, since they were recorded for the soluble monomeric FapC, however shown here at the structural model representing the amyloid fibrillar conformation.

### FapA is an intrinsically disordered chaperone and interacts with FapC

FapA is one of the six proteins expressed within the functional amyloid formation related fap operon (fapA-F) of *Pseudomonas aeruginosa*, and proposed to be a chaperone.^42^ FapA is an IDP similar to FapC based on the 2D HSQC fingerprint spectrum of the ^15^N labeled FapA and small chemical shift dispersion on the proton axis, **Figure 1C**. We presented the complete NMR resonance assignment of FapA recently (BMRB #52035) and the SSPs are shown in **Figure 3A**,^43^ along with the secondary structural elements from the Alphafold model (uniport #C4IN68) in **Figure 3A,B**. Recently it was shown that IDP FapA affects FapC fibrilization monitored by ThT assay, through an NMR-invisible oligomeric species which did not affect the NMR spectrum of FapC.^17^ Since one of the major differences between the functional and pathologic amyloids is the tight control of functional amyloid formation, the FapA chaperone is a candidate to interact and therefore regulate FapC.^42^ This mechanism was proposed to prevent self-harm to bacteria by eliminating undesired off-pathway fibril formation.^6, 44^ Conditional disorder of a chaperone for its action is known, as well as IDPs acting as a shield thereby slowing down fibrillation.^45, 46^ Non-chaperone IPD proteins were shown to slow down the aggregation of amyloids, such as Aβ by interfering with the nucleation phase, which can be correlated to FapA interaction with FapC.^46^ However, fully IDP protein acting as a chaperone and having significant effect on the monomeric prefibrillar species has not been reported previously in the literature for other systems.

**Figure 3.**
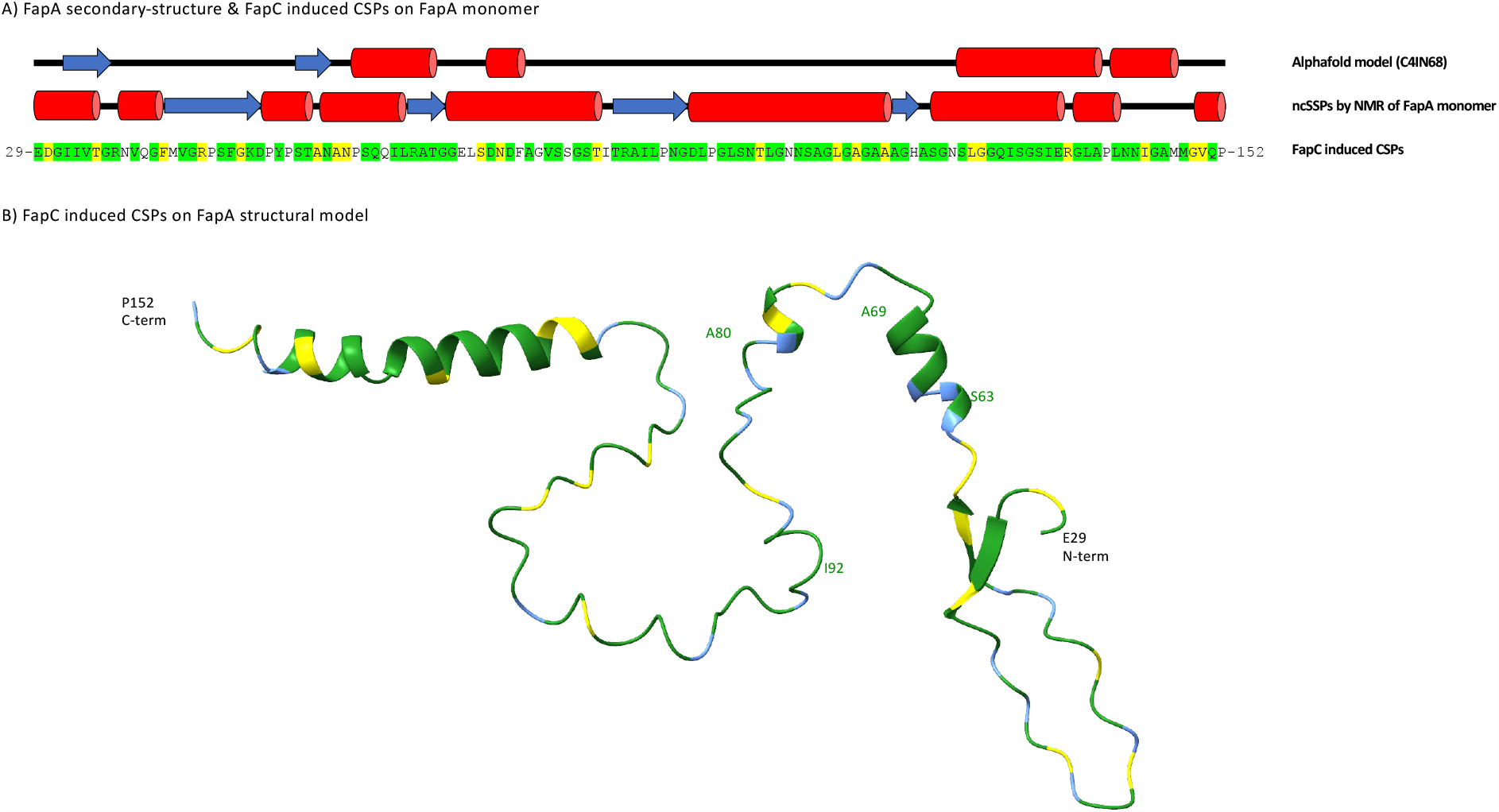
**A)** The secondary structure comparison of the FapA structure based on Alphafold model (AF-C4IN68) and the neighbor-corrected secondary structure propensities (ncSSPs) based on our experimental solution NMR chemical shifts of the monomeric FapA with the BMRB entry #52035. The amino acid sequence of FapA (29-152 residues) are shown at the bottom. The Alphafold secondary structure elements are depicted by the model. We used a threshold of 3-residues for depicting the α-helices (red cylinders) or β-sheets (blue arrows) as a secondary structure element for the NMR ncSSPs data. The yellow (medium CSPs) and green (large CSPs) highlights at the aminoacid sequence are following the notation described in Figure 1F, and the CSPs shown in Figure 1G. The FapA sequence is shown to visualize the CSPs, and the yellow highlight indicates significant changes, and the green highlight indicates much larger changes as depicted with two dashed lines in the CSPs plot in Figure 1G. **B)** Representation of the FapC induced shifts on FapA Alphafold structural model. The CSPs are shown on the structural model that are observed in the solution NMR spectra of isotope labeled FapA as a soluble IDP upon addition of four-fold unlabeled FapC monomeric protein.

Even though it is not straightforward to think of an IDP acting as a chaperone, by using high-resolution solution NMR data we clearly show here that IDP FapA acts as a chaperone (or chaperone-like) for monomeric FapC *in vitro*, interacting with prefibrillar monomeric FapC and slows-down FapC fibrillation through specific interactions, **Figure 1D**,**G** and **3**. We monitored FapA/FapC interaction from both FapC and FapA perspective, when they are complex with four-fold access corresponding partners, FapA or FapC, respectively. The NMR interaction experiments monitored from the FapC perspective utilizing ^15^N isotope-labeled FapC and unlabeled FapA sample and from the FapA perspective utilizing ^15^N isotope-labeled FapA and unlabeled FapC sample. 2D ^1^H-^15^N HSQC NMR spectra of the pure FapC and FapC:FapA samples were recorded under identical conditions and shown in **Figure 1A-B**. The 1-day incubation of FapC with excess FapA induces NMR spectral changes through chemical shift perturbations (**Figure 1B,F** and **2A**) and signal intensity changes (**Figure 4F**).

**Figure 4.**
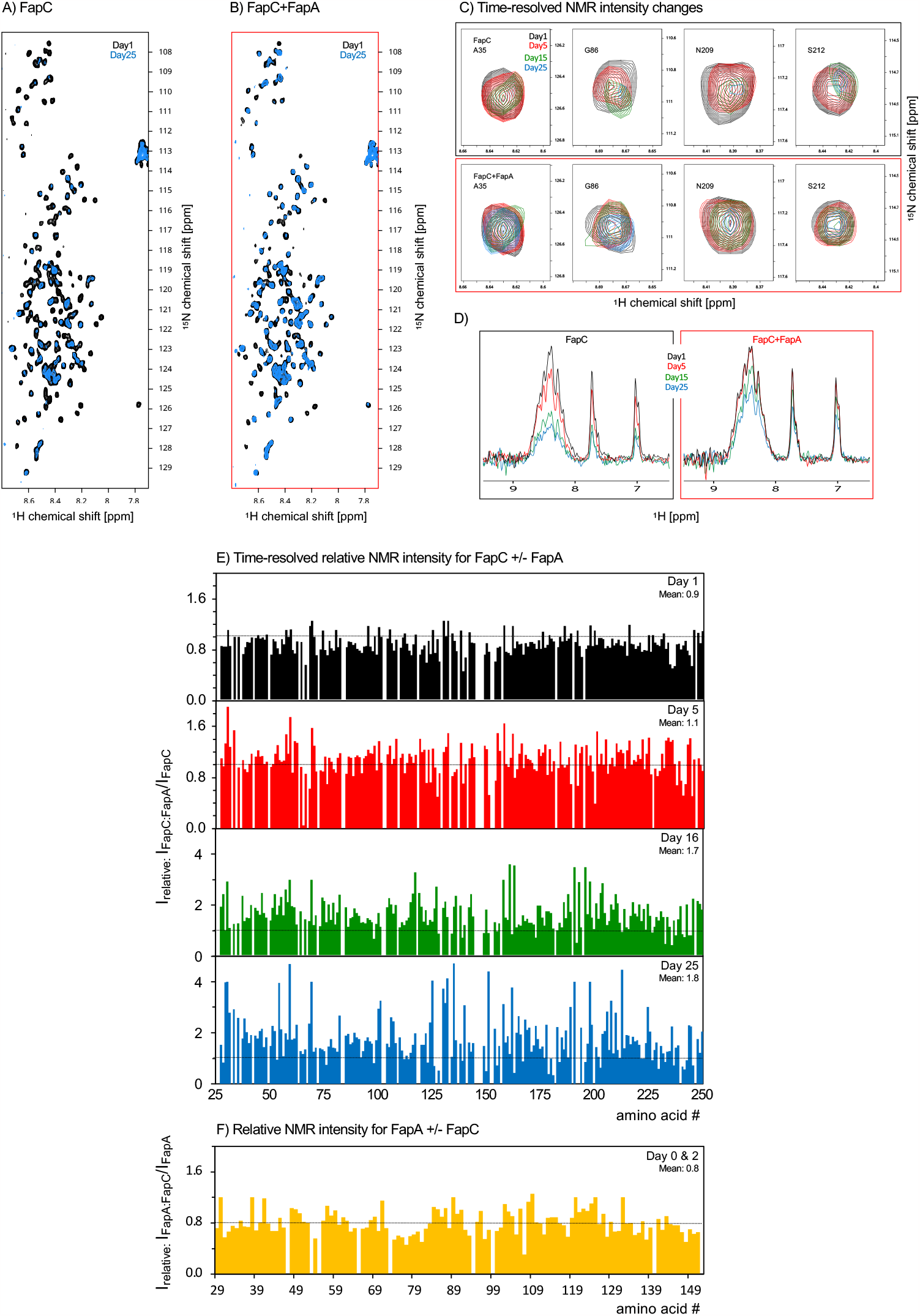
Time-dependent 2D ^1^H-^15^N HSQC NMR spectra for pure **A)** FapC and **B)** FapC:FapA 1:4 samples. The spectra were recorded over 25 days at Day 1, 5, 16 and 25 for both samples under identical conditions and plotted with the same contour levels for direct comparison. **C)** Zoomed regions for selected amino acids are shown for pure FapC (top) and FapC:FapA 1:4 (bottom), with the same color coding as in A/B. **D)** The first spectrum of each 2D ^1^H-^15^N HSQC spectra is used to quantify the signal amount plotted here by signal integration at the amide proton region. **E)** Representation of the residue-resolved NMR signal intensity ratios for FapC:FapA samples at four-fold excess mole ratio of the unlabeled FapA. The signal intensity of the NMR signal from each residue of the FapC:FapA 1:4 sample is divided with the signal intensity of the same residue of the pure FapC sample. These plots show an estimate of the time-dependent effect of FapA chaperone on the overall signal amount observed in FapC NMR spectrum at 1, 5, 16 and 25 days of incubation time. It is a residue-specific representation of the bulk signal intensity decay results shown in A and B. The average values of the relative intensities are given at each plot, as well as the dashed line at y-value=1. Overall, it is clear that FapA significantly prevents the FapC signal intensity decrease due to fibrillation compared to the pure FapC. Note different scales on the y-axes. **F)** Representation of the residue-resolved NMR signal intensity ratios for FapA:FapC samples at four-fold excess mole ratio of unlabeled FapC, at both 0/2 days of incubation time where incubation longer did not change the intensity ratio.

The FapA induced chemical shift perturbations (CSPs) on FapC NMR spectrum are detectable and significant as shown by shifts exceeding the two different confidence levels (CSPs greater than “mean + standard-deviation” or “mean + 2*standard-deviation”), **Figure 1F**. Previous interaction studies on IDP proteins suggest chemical shift changes of similar magnitude as significant shifts, e.g. on CsgA and αSynuclein protein.^15, 47^ Remarkably, much greater CSPs were detected on the FapA 2D NMR spectrum when the interaction experiment is done under identical conditions from the FapA perspective by incubating ^15^N labeled FapA with four-fold access unlabeled FapC, **Figure 1C,D**. The CSPs plot in **Figure 1G** represents up to six-fold larger perturbations compared to the FapA induced perturbations in FapC spectrum. The average weighed cumulative CSPs are 0.015 ppm and 0.087 ppm for FapC:FapA (**Figure 1B,F**) and FapA:FapC (**Figure 1D,G**), respectively. Overall, 41 out of 226 residues (∼18%) show significant CSPs spread over the whole FapC sequence upon addition of FapA. Remarkably, 110 out of 123 residues (∼90%) show large CSPs on FapA spectrum upon addition of FapC spread over the whole sequence, **Figure 1G**, determined by using the significance levels in **Figure 1F**. These CSPs are shown on the FapA Alphafold structural model in **Figure 3A,B**. The four largest CSPs are observed for residues S63, A69, A80 and I92. In summary, these two different NMR experimental setups from different aspects of FapC-FapA complex clearly show that FapC and FapA are interacting.

### FapC and FapA interaction interface by solution NMR CSPs

The residue-specific FapA induced CSPs in the ^15^N labelled FapC 2D NMR spectrum are visualized on the FapC amyloid structural model based on Alphafold’s prediction and our NMR resonance assignment, **Figure 1F** and **2A**,**D**.^48^ Strikingly, the largest changes with values greater than the “mean + 2*standard-deviation” out of the significant CSPs, are observed predominantly at eleven residues at A82, L158, I160, A169, H191, S192, N201, C237, C240, A241 and G245. These could indicate the hotspots of fibrillation inhibition/control interface between FapA and FapC. Particularly, the A82, N201, C237, C240, A241 and G245 have the largest CSPs, and this may indicate a more specific interaction of FapA at the R1/L1 and L2/R3 transition area, and at the C-terminus. Previous ThT assays performed on R3C, L2R3C, L1L2R3C and FL FapC constructs indicate that the L1 and L2 loop regions increase the aggregation propensity without changing the fibril morphology,^37^ can be correlated to the largest CPSs. **Figure 2A**,**D** represents the location of these residues showing large CSPs at the structural model and surprisingly many of them are clustered close to each other at the C-terminus and linkers between the β-sheets. It should be noted that the model represents the structure of the folded fibrillar FapC, whereas the CSPs were observed for the monomeric prefibrillar IDP FapC having predominantly random coil conformation with mentioned sizable SSPs. Nevertheless, the localization of CSPs may suggest hot spots for chaperone-based prevention/slowing-down of functional amyloid formation. The electrostatic potential of the structural model shown in **Figure 2C** suggest that the CSPs are not at the most charged areas, supporting the FapA and FapC interaction to be structurally specific. The signal intensity changes observed in the 2D NMR spectrum of FapC upon addition of FapA at 1:4 mole ratio for samples supports the interaction of FapA with FapC, see **Figure 4E**.

To further support the FapC and FapA interaction, we also performed the FapC and FapA interaction experiment from the point of ^15^N labeled FapA and monitored changes in chemical shifts upon the addition of unlabeled FapC, **Figure 1C**,**D** and **G**. Remarkably, we observed CSPs six-fold larger compared to the FapA induced changes in the FapC spectrum. This clearly supports that FapA and FapC interacts with each other as IDP monomers, which is observable at our experimental condition of 274 K and pH 7.8 with samples close to the native amino acid sequence without any large tag. ∼90% of FapA residues represent strong CSPs, **Figure 1G** and **3A**,**B**.

### FapA strongly decelerates FapC fibrillation

To quantify the time-resolved effect of FapA on FapC fibrillation, we recorded 2D ^1^H-^15^N HSQC spectra of pure FapC and FapC:FapA 1:4 samples, **Figure 4A**,**B**. All the spectra were recorded at 274 K and the samples were stored at 4 °C in between the NMR measurements to keep the same conditions throughout the experimental time span of 25 days. Lowered temperatures allowed first, to observe all the signals in FapC and FapC:FapA samples at the current sample conditions and second, more importantly to slow down the fibrillation so that we can monitor changes correctly without any transient effects. Temperature dependent slowing down of fibrillation were reported similarly before by using ThT assays.^32^ This simple yet powerful experiment can reveal the residue-specific aggregation propensities by quantifying the residue-resolved NMR intensity losses over time, **Figure 4E**. One can think of this experiment as a ThT assay that can monitor changes at the residue level in the protein. The transition of the soluble FapC monomer into high molecular-weight species (soluble/insoluble oligomer/fibril) results in solution NMR signal intensity attenuation, since solution NMR predominantly monitors the low molecular weight species, i.e., monomeric FapC.^49^

To determine the time-dependent NMR signal loss, first a qualitative analysis is performed by the integration of the protein backbone amide signal intensities of the first slices of each 2D HSQC experiments, **Figure 4D** and **5C,D**. This data shows that FapA slows down the bulk FapC signal intensity decay at the backbone amide region. Over 25 days, the 1D NMR signal for pure FapC and FapC:FapA samples are reduced by ∼69% and 34%, respectively. This observed FapA-aided slowing down in fibrillization strongly supports the interaction of FapC and FapA. Our data correlate with the similar 1D NMR signal intensity monitoring of amyloid fibril fibrillation previously reported on pathological Aβ and Huntington fibrils as well.^26, 29^

The overlay of the 2D HSQC recorded at 1 and 25 days of incubation for FapC and FapC:FapA samples are shown in **Figure 4A,B**. The reduction in the overall intensity and number of observed peaks demonstrate that the soluble monomer to high molecular weight species transition occurs slowly at 274 K. From these 2D spectra, residue-specific quantitative signal intensity reduction analysis is performed for the spectra recorded at 1 (black), 5 (red), 15 (green) and 25 (blue) days of incubation at 274 K, **Figure 4E**. Additionally, representative NMR signals from the A35, G86, N209 and S212 are shown in **Figure 4C** as zoomed regions for visualization of the time-dependent signal reduction for FapC and FapC:FapA 1:4 samples.

At 1 day of incubation time the FapC:FapA sample has a slightly reduced NMR signal intensity compared to the pure FapC with an average relative ratio ∼0.9 (I_relative_: I_FapC:FapA_ / I_FapC_), **Figure 4E** (top raw, black). This FapA induced NMR signal intensity reduction supports that FapA is interacting with FapC as indicated by the CSPs, **Figure 1A,B**. The NMR measurements performed at longer incubation times at 274 K (at 5, 16 and 25 days) show that FapA containing FapC has much larger NMR signal intensity compared to the pure FapC up to a factor of ∼2. At day 5, I_relative_ is 1.1, and further increases to 1.7 and 1.8 at day 16 and 25, respectively. FapA keeps FapC monomeric to a higher degree by slowing its fibrillation, resembling the experiments done on CsgA functional amyloid from *E. coli* incubated with its CsgC and CsgE chaperone partner.^15, 50^ Increased NMR signal intensity was observed compared to the pure CsgA sample, upon incubation with CsgE chaperone for one day at 277 K and measured at 284 K. Surprisingly, the time-resolved intensity ratio data shown in **Figure 4E** are uniform with similar overall signal intensity ratios at day 1 and 5, however, becomes less uniform and show deviation from unity for later time points recorded at day 16 and 25. The emerging differential I_relative_ pattern is not easy-to-understand, nevertheless, we speculate that signals from the N-terminus and R1-R2-L2-R3 regions maybe differentiating compared to the rest of the sequence at longer incubation times, to be further investigated.

### FapA – FapC interaction and chaperone effect is temperature dependent

We have performed time-dependent solution-state 1D NMR and ThT fibrillization experiments at different experimental temperatures between 274 and 310 K for pure FapC, pure FapA and FapC:FapA samples (at 1:4 and 1:10 mole ratio). This was to quantify the effect of temperature on the FapC fibrillation as well as the effect of FapA on this process, since temperature remains a parameter that is different between this study and previous work on FapC-FapA system. We freshly desalted FapC and prepared corresponding pure FapC and FapC:FapA 1:4 samples for the NMR and ThT experiments. We used 50 μM FapC, 50 μM FapA and corresponding concentrations of FapA for the complexes (1:4 and 1:10). As an established method, ThT assay monitors the amyloid fibril formation, but do not respond to non-filamentous prefibrillar species or amorphous aggregates, **Figure 5A**,**B**. On the other hand, solution NMR can monitor the soluble monomeric FapC and the signals from the high-molecular weight fibrillar or oligomeric species will disappear from the spectra, **Figure 5C**,**D**. At t:0, the samples are monomeric, and the maximum amount of solution NMR signal is observed, whereas the ThT response is zero since no fibrils formed yet. Over time, NMR signal intensity decreases and the ThT signal increases due to the transition of FapC from monomer to fibril. We did not observe a lag-phase for the pure FapC and FapC:FapA samples, especially at 310 K, nevertheless, overall, the ThT data is similar to previous data.^44^ 310 K is a common temperature for ThT fibrillization monitoring assay,^44^ and to correlate the ThT results to the NMR results recorded at lower temperatures, we performed the ThT fibrilization assay at 277 K and monitored intermittently over 24 hours. From the ThT perspective, experiments performed at 310 K indicate that FapA slows down the fibrilization kinetics in a concentration-dependent manner and reduces the end point fluorescens, **Figure 5A**. As a control, the pure FapA was measured which did not give any ThT response. At lower temperature of 277 K, the effect of FapA on the FapC fibrilization is reduced, nevertheless still present, **Figure 5B**. Slowing down of fibrillization kinetics at lower temperatures results in different shapes in the ThT curves.

**Figure 5.**
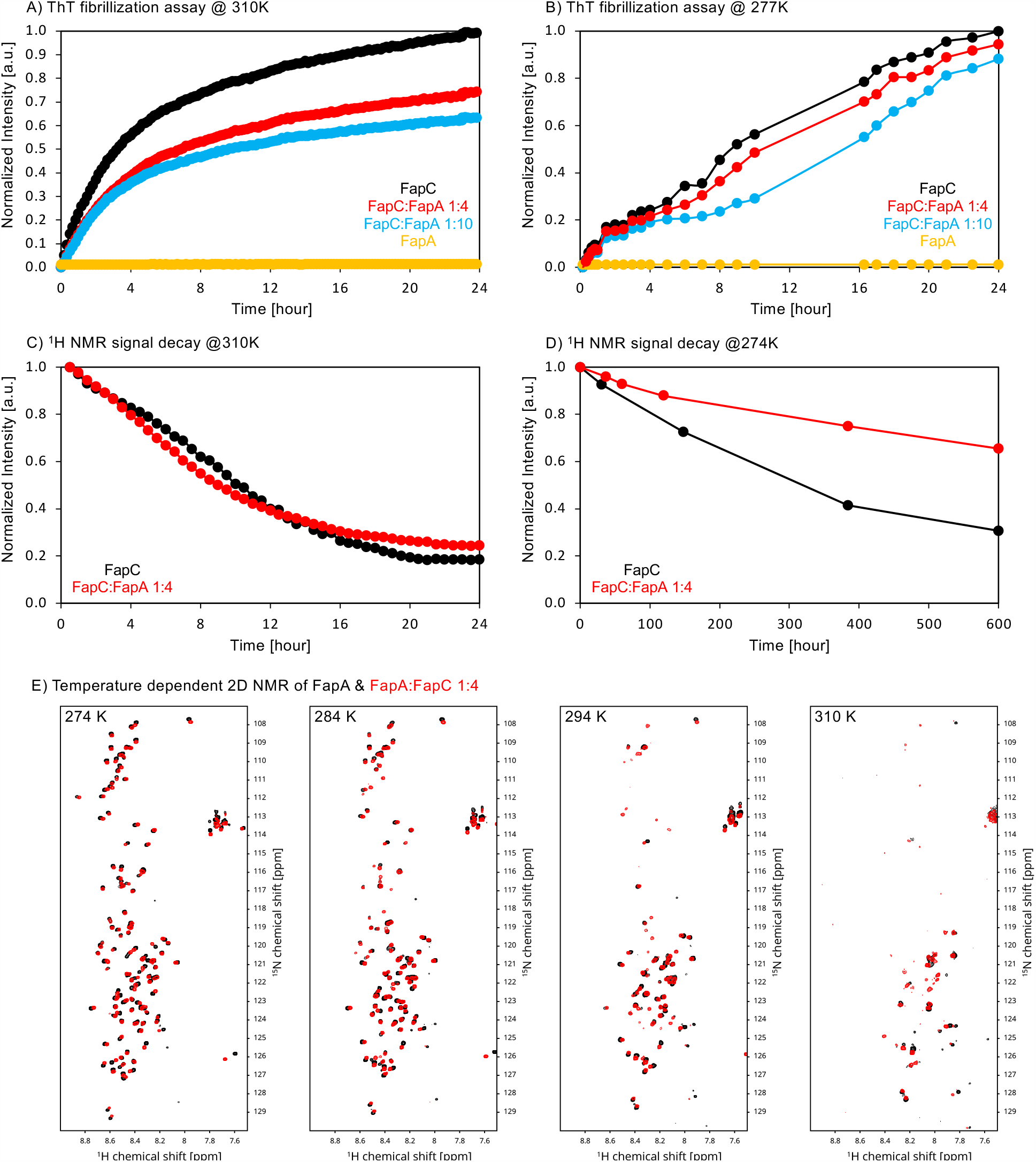
The temperature dependent FapA – FapC interaction monitored by ThT fibrillation assay and ^1^H NMR. **A**,**B)** ThT fibrillization assay to monitor the monomer to amyloid fibril transition at 310 and 277 K. 50 μM FapC (black), FapC:FapA1:4 (red), FapC:FapA1:10 (blue), 50 μM FapA (yellow). **C**,**D)** Time-dependent ^1^H NMR signal intensity reduction for pure FapC (black) and FapC:FapA1:4 (red) samples at 310 and 274 K. **E)** The 2D ^1^H-^15^N 2D HSQC spectra recorded for FapA and FapA:FapC 1:4 samples at four different temperatures, to show the effect of temperature to the CSPs observed at ^15^N isotope-labeled FapA upon the addition of four-fold excess unlabeled FapC. The CSPs persists at all the NMR experiment temperatures, although higher temperatures result in signal loss for many resonances due to increased amide proton exchange. The spectra were referenced at each condition by using internal reference DSS.

From the NMR perspective, at 274 K four-fold excess FapA significantly reduces the NMR signal intensity loss extent over the 600 hours of total experimental monitoring time compared to the pure FapC, **Figure 5D**. Due to the low temperature, the fibrillation kinetics is much slower and the NMR monitoring experiment was run much longer than the regular 24-hour time span.^32^ However, temperature strongly changes the chaperonic effect of FapA on FapC, and we observed that the fibrilization deceleration effect of FapA is abolished in the NMR monitoring experiment to a great extend at 310 K, **Figure 5C**, and the FapC and FapC:FapA NMR signal decay curves become similar with small differences at the overall shape and end point intensity. The 2D HSQC spectra recorded between 274 and 310 K, **Figure 1D** and **5E**, indicates that the CSPs persist at all the temperatures for the ^15^N isotope-labeled FapA upon the addition of four-fold excess unlabeled FapC, possibly with smaller magnitude at higher temperatures since the perturbations for some residues became smaller. More quantitative analysis was not performed due to the signal loss at higher temperature due to increased amide proton exchange.

Overall, these ThT and NMR data indicates that the chaperone effect of FapA on FapC is temperature dependent. This could be the reason of the discrepancy between the previously reported FapC and FapA interaction studies and results shown here. Surprisingly, the ThT and NMR data do not align perfectly, indicating differences in what types of protein species these techniques monitor.

### FapA does not alter FapC fibril morphology

Negative-staining transmission electron microscopy (TEM) was performed to identify the morphology of the fibrils formed by FapC samples with and without FapA. The NMR samples are used right after the 25 days of NMR measurement time cycle to prepare the EM samples and to record the micrographs shown in **Figure 6** for the pure FapC and FapC:FapA 1:4 samples. Overall, the pure FapC sample shows very high quality throughout the micrographs, with long and mostly single filaments with an averaged diameter of 7.8 ± 1.3 nm, determined from ∼200 measurements on different filaments from different EM grids, **Figure 6A**,**C** and consistent with previous reports.^17, 37^ Small amount of amorphous aggregates were found for the pure FapC sample in the grids and observed consistently for different EM grid preparations.

**Figure 6.**
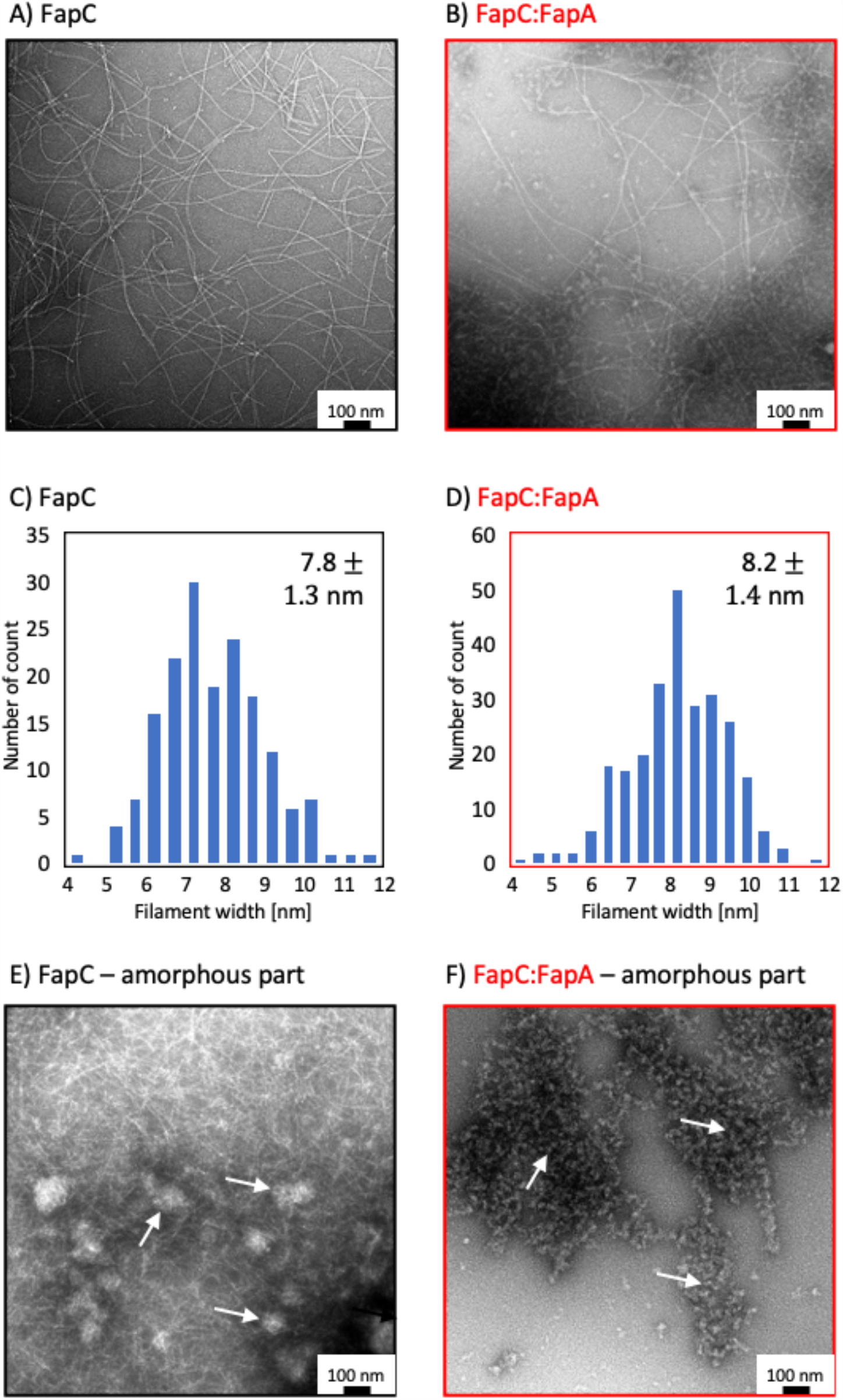
The negative-staining transmission electron microscopy (TEM) images of the fibrils of **A)** pure FapC and **B)** FapC:FapA 1:4 samples. The averages of ∼200 filament width measurements are shown as histograms for **C)** pure FapC and **D)** FapC:FapA 1:4. Similar filament widths are observed for both samples. The FapC contains predominantly pure filaments with a small fraction of amorphous fraction. Larger amorphous aggregate fraction is observed for FapC:FapA 1:4 sample along with still high-quality filaments. The amorphous parts are shown in **E)** for pure FapC and **F)** for FapC:FapA 1:4 samples, respectively, indicated with white arrows.

The FapC:FapA 1:4 sample also contains long filaments similar to the ones observed for FapC with an averaged filament diameter of 8.2 ± 1.4 nm, **Figure 6B**,**D**. The filament widths for the pure FapC and FapC:FapA samples are similar within the experimental error. However, the micrographs prepared from the FapC:FapA 1:4 sample contain a much larger amount of amorphous aggregates as standalone moieties or attached to the FapC filaments (see **Figure 6E**,**F**), which was similarly observed previously.^17^ We note that the amorphous aggregates have different appearance and ratio for both samples, and are due to FapC in the pure FapC sample, but most probably due to excess FapA in the FapC:FapA sample. The amorphous aggregates observed for FapC:FapA sample shown in **Figure 6F** were not detected for pure FapC sample, and same type of aggregates were observed for the micrographs prepared by pure FapA (data not shown).

## Discussion

We employed high-resolution solution NMR spectroscopy to investigate the prefibrillar monomeric form of the biofilm-forming FuBA FapC protein from *Pseudomonas*. By analyzing its secondary structural propensity, we found that FapC is primarily in a random coil or PP2 helix conformation, with notable α-helical and β-sheet propensities. The PP2 and the pre-structural α-helical and β-sheet propensities are key towards structural transition to the correct amyloid fibrillar fold. These secondary structure propensities, along with the linker regions separating them, align well with the secondary structure motifs on the Alphafold FapC amyloid structure model. Moreover, the *in silico* predicted APRs fall within these β-sheet predicted regions in the structure. Two α-helical regions indicated by the ncSSPs do not align with the structural model or APR predictions, highlighting disparities between the NMR-determined intrinsically disordered protein (IDP) structural elements of FapC and its amyloid fold. Overall, the agreement between the IDP secondary structure propensities and the predicted model structure suggests a smooth/unhindered transition towards the amyloid fold without large structural rearrangements.

We examined the influence of the FapA chaperone on the fibrillation of FapC. FapA is a component of the functional amyloid operon that regulates FapC *in vivo*, preventing undesired fibrillation.^9^ Our solution NMR studies revealed that, similar to FapC, FapA is also an IDP. Time-dependent NMR spectroscopy and ThT analysis provided direct evidence that FapA acts in a chaperone-like manner, significantly decelerating FapC fibrillation, making this the first reported instance of an IDP with chaperone activity. Consequently, we observed approximately a twofold increase in the amount of soluble monomeric FapC through time-resolved 1D and 2D NMR experiments. Upon incubation of ^15^N labeled FapC with unlabeled FapA, we observed CSPs in the FapC spectrum, as well as intensity changes for the FapC:FapA samples compared to FapC. These CSPs were not previously detected by NMR.^17^ Notably, 36 residues displayed significant shifts, primarily concentrated in specific regions predicted by the FapC fibrillar structural model. Among these residues, N201, C240, A241, and G245 exhibited the largest CSPs, indicating a potential interaction of FapA with FapC at the crossing interface of L2 and R3, as well as at the C-terminus. Furthermore, counter NMR titration experiments, monitoring the effect of unlabeled FapC on the ^15^N labeled FapA NMR spectrum, demonstrated CSPs approximately six times larger, with significant shifts observed in ∼90% of the NMR signals. Among these, S63, A69, A80 and I92 have the largest CSPs which are clustering around the middle of FapA. Collectively, these findings strongly suggest an interaction between the monomeric IDP proteins FapA and FapC, motivating further comprehensive experiments to quantitatively assess the structural mechanisms underlying chaperone-assisted amyloid interactions and prevention within the fibrillar FapC fold.

We conducted time-resolved NMR measurements to track changes in signal intensity for pure FapC and FapC:FapA samples at a 1:4 ratio, providing further evidence of the interaction between the FapA chaperone and FapC fibrillation over a 25-day experimental period. The use of lower NMR measurement temperatures was crucial for accurate and timely observation of these changes, enabling the monitoring of time-dependent resonance intensity alterations. At day 1 of incubation, the FapC:FapA sample exhibited an overall reduction in signal intensity compared to pure FapC, indicating the interaction between FapA and FapC. Similarly, when FapC was added to the FapA, FapA NMR signal intensity is reduced to ∼80% at day 0 and 2 incubation times. Subsequently, from day 5 to day 25 of incubation, the average intensity of the FapC samples incubated with FapA increased nearly twofold compared to pure FapC. These results collectively demonstrate the chaperone effect of the intrinsically disordered protein FapA on the functional amyloid FapC. FapA not only interacts with monomeric FapC but also slows down its fibrillation. Interestingly, FapA displays chaperone-like properties similar to the well-folded CsgE nonameric chaperone from the *E. coli* functional amyloid system, despite distinct structural characteristics of these two proteins as being folded versus unfolded. Furthermore, we have identified residue-specific interaction hotspots that provide insights into the structural basis of chaperone-mediated prevention of functional amyloids. The residue-specific NMR resonance intensity ratios for pure FapC and FapC:FapA 1:4 samples exhibited uniformity at 1 to 5 days of incubation. However, at day 16 and 25, variations among the sequence were observed, indicating potential differential effects of FapA on FapC over time. Specifically, the N-terminus and R1,L2,R3 regions displayed higher overall NMR resonance intensities compared to the rest of the FapC sequence, suggesting a more specific and/or preferential action of FapA on these regions. More remarkably, when FapC was added to the ^15^N labeled FapA, ∼90% of the signals showed large CSPs spread over the whole FapA sequence.

The temperature dependent ThT and NMR signal decay analysis give insights into the fibrillization kinetics as a function of temperature. At the conventional 310 K measurements, the reduction in the end point fluorescens value at the FapC ThT response as a function of FapA concentration is observed. This indicates that FapA reduces FapC fibrillation and supports the chaperonic properties of FapA. Moreover, the same ThT experiments performed at 274 K, indicates a similar effect of FapA on FapC fibrillization. The ^1^H NMR measurements performed at these temperatures to quantify the reduction in the amount of soluble monomer by recording the NMR signals, indicates additional insights. 274 K NMR data resembles the ThT data and indicates FapA reducing FapC fibrillization. However, the 310 K NMR data shows only a minor change in the signal decay properties and indicates that the NMR observable chaperonic effect is abolished. Overall, these data suggest a temperature dependent chaperonic FapA effect on FapC, and differences between the ThT and NMR based quantification.

TEM analysis of fibrillar samples revealed similar filament widths of approximately 8 nm, indicating that FapA does not alter the FapC filament morphology. However, a significant difference was observed in the amorphous fraction of the samples on the EM grids. The FapC:FapA sample exhibited a larger amount of amorphous material observed throughout the grids, which also seemed to stick to FapC filaments. While FapC also formed a small amount of amorphous aggregates, the greater abundance of such aggregates in the FapC:FapA sample was likely due to the excess FapA that precipitated during grid preparation.

In conclusion, our study employed solution-state NMR spectroscopy to investigate the functional amyloid FapC at a residue-level resolution. Through the analysis of high-resolution 2D ^1^H-^15^N HSQC NMR spectra, we obtained precise information regarding the secondary structure elements of FapC, which exhibited a close alignment with the predicted fibrillar structural model generated by Alphafold. Our findings quantitatively characterized the impact of the intrinsically disordered chaperone FapA on FapC fibrillation, providing clear evidence of their interaction and the ability of FapA to decelerate fibrillation kinetics. Notably, FapA did not induce any discernible alteration in the final morphology of FapC filaments, as confirmed by electron microscopy observations. Furthermore, our study offered valuable insights into the structural aspects of the FapC-FapA interaction, establishing correlations with the amyloid predicted structural model. Based on our comprehensive findings, we propose a model in which FapA achieves its fibrillation-slowing effects by specifically engaging with monomeric FapC at the flexible loops and C-terminus, as supported by the spatial clustering of CSPs on the Alphafold structural model.

## Materials and Methods

### Recombinant protein production

Details of FapC protein expression and purification is described in detail previously,^31^ we give a short summary here. The full-length FapC (residues 25-250, 226 aa) was cloned into a pET28a vector system. There are nine additional amino acids in the produced protein samples due to the construct design, one methionine on the N-terminus and a histidine purification tag of LEHHHHHH at the C-terminus. Bacterial plasmids were transformed into the BL21(DE3) *E. coli* bacteria. Minimal media containing ^15^N-ammonium chloride is used to uniformly ^15^N isotope label FapC. The transformed BL21(DE3) cells were plated onto LB agar and grown overnight in 37 °C incubator. The colonies were resuspended and inoculated into large volume media. These cultures were grown in shaker incubator at 37 °C until the OD600 reached a value between 0.8-1.0 OD. For induction, 1 mM IPTG was added to the culture and bacteria was grown for 4 hours at 37 °C. Cells were harvested by 6000 RCF centrifugation for 20 min at 4 °C. The cells were lysed using a sonicator in denaturing buffer. His-tag affinity column was used to purify the protein by step-eluting the protein. FapC containing fractions were pooled together and spun down by 20800 RCF, 10 min at RT. The supernatant was stored at -80 °C.

FapA gene block was ordered from Twist Bioscience and primers (IDT DNA) were used to clone into pET32 vector using In-fusion cloning mechanism (Takara Bio). The FapA construct without the signal sequence (residues 29-152, 124aa) was expressed with an N-terminal Thioredoxin (Trx) followed by a 10 histidine (His10) tag and thrombin cleavage site. Due to the lack of a tryptophan, one was engineered into the C-terminal end of the protein for accurate protein quantification. The final cleaved FapA protein contains five additional amino acids, four at the N-terminus as cloning and cleavage artifacts (GSGT) and one at the C-terminus (W).

FapA plasmid was transformed into BL21(DE3) e. coli bacteria. The transformed BL21(DE3) cells were plated onto LB agar with ampicillin and grown overnight in 37 °C incubator. The colonies were resuspended in LB media with ampicillin antibiotic and inoculated into large volume media. These cultures were grown in shaker incubator at 37 °C until the OD600 reached a value between 0.8-1.0 OD. For induction, IPTG was added to each culture for a final concentration of 1 mM and grown for another 4 hours at 37 °C. Cells were harvested by centrifugation (6000 RCF, for 20 min at 4 °C). Cells were resuspended in resuspension buffer (50 mM Tris, 300 mM NaCl, 2 mM PMSF, pH 8.0) and lysed using a sonicator. Cell lysate was centrifuged (29,000 RCF, for 20 min at 4 °C), and the soluble fraction was decanted and collected. Trx-FapA was purified from the soluble fraction using a His-Tag affinity column using a gradient elution from 0-500mM imidazole (in addition to 50 mM Tris, 300 mM NaCl, 2 mM PMSF, pH 8.0). The eluted Trx-FapA was pooled and dialyzed into 20 mM Tris pH 8.7 at 4 °C. Thrombin was added to the dialysis bag for the last dialysis at 4U per mg of Trx-FapA, and cleavage was done overnight at 4 °C. The cleaved FapA was purified using a SourceQ column over a 0-100 mM NaCl gradient (with 20 mM Tris, 2 mM PMSF, pH 8.7). Pure FapA was pooled and stored at -80 °C.

### NMR Spectroscopy

2D ^1^H-^15^N HSQC NMR spectra of uniformly ^15^N isotope labeled FapC FL were recorded at a Bruker Avance III 600 spectrometers equipped with a 5 mm triple resonance TCI cryoprobes. 3 mm NMR sample tubes were used. A total volume of 550 μl of 50 μM protein was prepared in 20 mM sodium phosphate, 1 mM D6-DSS, 10% D_2_O, 0.02% Sodium Azide (w/v) and at pH 7.8. The NMR samples were prepared freshly prior to performing the NMR experiment. All the 2D ^1^H-^15^N HSQC NMR experiments were recorded at 274 K. A total of 32 scans were recorded for 300 increments, with 0.9 s recycle delay. We previously described the details of the FapC and FapA NMR resonance assignment, and the data were deposited to BMRB (FapC: 51792/51793 and FapA: 52035).^31^ The ^1^H chemical shifts were referenced at each temperature directly to 0 ppm by using DSS as an internal standard, and the ^15^N chemical shifts were indirectly referenced by using the ^1^H frequency.^51^ 2D ^1^H-^15^N HSQC NMR spectrum of uniformly ^15^N isotope labeled FapA (29-152 amino acids, without the signal peptide). The NMR spectrum was recorded at a Bruker Avance III 600 spectrometer equipped with a 5 mm triple resonance TCI cryoprobes at 274 K. 5 mm NMR sample tube was used. A total volume of 550 μl ∼0.34 mM protein was prepared in 20 mM sodium phosphate, 1 mM D6-DSS, 10% D_2_O, 0.02% Sodium Azide (w/v) and at pH 7.8. The NMR sample was prepared freshly prior to performing the NMR experiment. A total of 32 scans were recorded for 300 increments, with 0.9 s recycle delay.

The FapC-FapA interaction assays were performed by preparing NMR samples with a constant concentration of ^15^N isotope labeled FapC at 50 μM, and the corresponding unlabeled FapA concentrations of 50 μM for 1:1 sample, 100 μM for 1:2 sample and 200 μM for 1:4 sample in 50 mM sodium phosphate, 30 mM DTT, 1 mM D6-DSS, 10% D_2_O and at pH 7.8.

The FapA-FapC interaction assays were performed by preparing NMR samples with a constant concentration of ^15^N isotope labeled FapA at 50 μM, and the corresponding unlabeled FapA concentrations of 200 μM for 1:4 sample in 50 mM sodium phosphate, 1 mM D6-DSS, 10% D_2_O and at pH 7.8.

### Negative-staining TEM Microscopy

400 mesh carbon film copper grids were glow discharged for 90 seconds. ∼2.5 μL of pure FapC and FapC:FapA 1:4 samples were applied to grids and left for 10 seconds before side blotting on filter paper. Grids were negative stained with 2% (weight/volume) uranyl acetate for 10 seconds before side blotting on filter paper. EM micrographs were recorded at a FEI Tecnai TF20 microscope with a field emission gun operating at 200 kV equipped with a TVIPS XF416 CMOS camera. No further image processing was done to the images shown.

### ThT assays

Tecan te-cool instrument was used for the ThT assays by using Corning 96-well plate (reference #3881) with half area. The polystyrene plates have non-binding surface and black clear bottom. The 37 °C (310 K) measurements were done by a controlled temperature at the instrument continuously. The 4 °C measurements were done by storing the plate at the fridge and measuring the ThT response every 30-60 minutes by with fast measurements (within 30 seconds) at an instrument equilibrated at room temperature. Afterwards, the plate was put back into the fridge and kept at 4 °C till the next measurement. This procedure was repeated for 24 hours.

## Data Availability

The datasets generated and/or analyzed during the current study are available in the BMRB repository, accession number FapC :#51792-51793 and FapA: #52035.

## Author Contribution

Conceptualization: UA; Methodology: CHB, MA, UA; Formal analysis and investigation: CHB, HS, KH, JJ, MA, UA; Writing -original draft preparation: CHB, MA, UA; Funding acquisition: MA, UA; Resources: MA, UA; Supervision: UA. All authors reviewed the manuscript.

## Acknowledgement

UA acknowledges financial support from University of Pittsburgh startup funding and the high-field NMR infrastructure at the Structural Biology Department, School of Medicine, University of Pittsburgh. MA acknowledges financial support from L’OREAL UNESCO For Women in Science. We acknowledge James Conway for help in the EM data collection.

## Supplementary Information

**Figure Supplementary 1:**
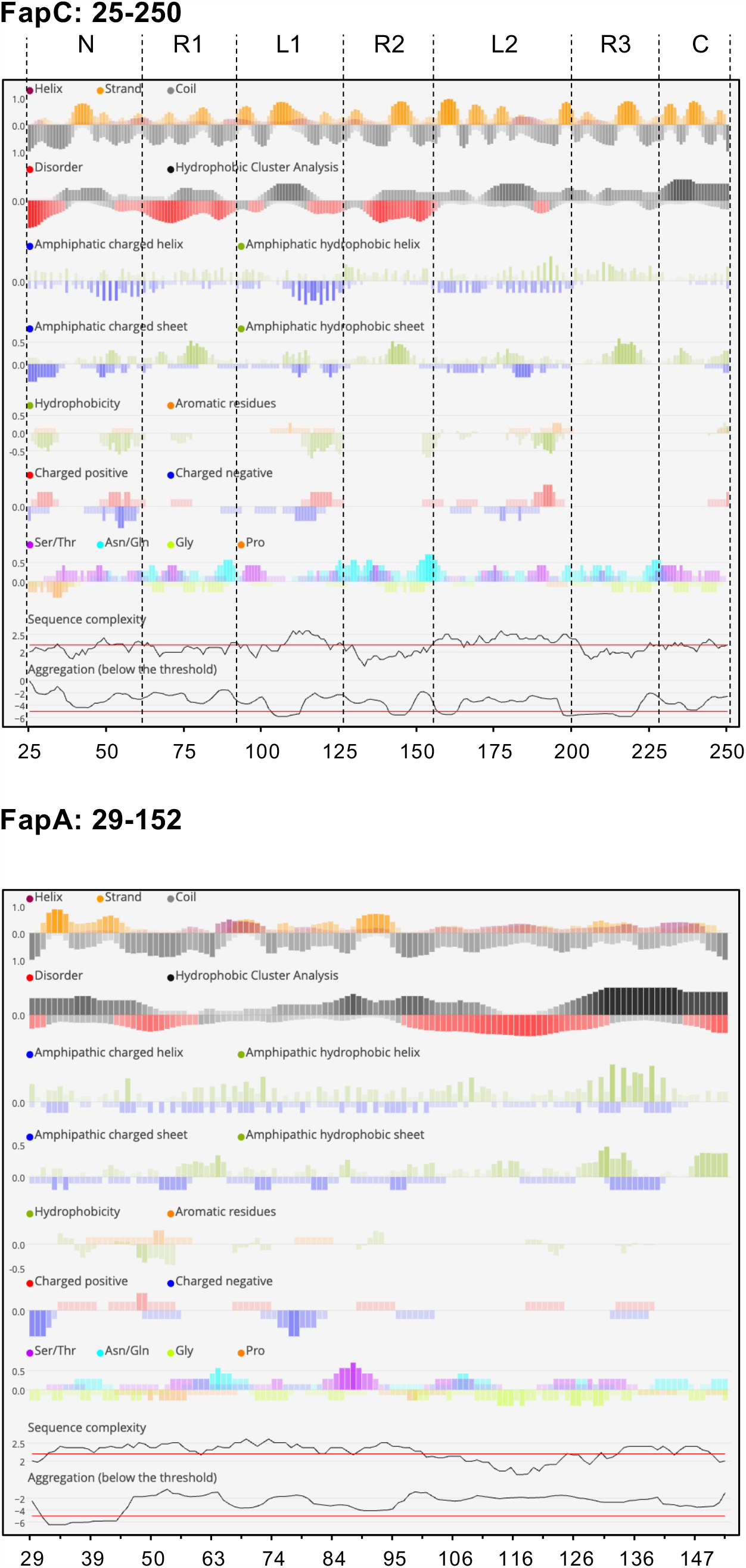

**Figure Supplementary 2:**
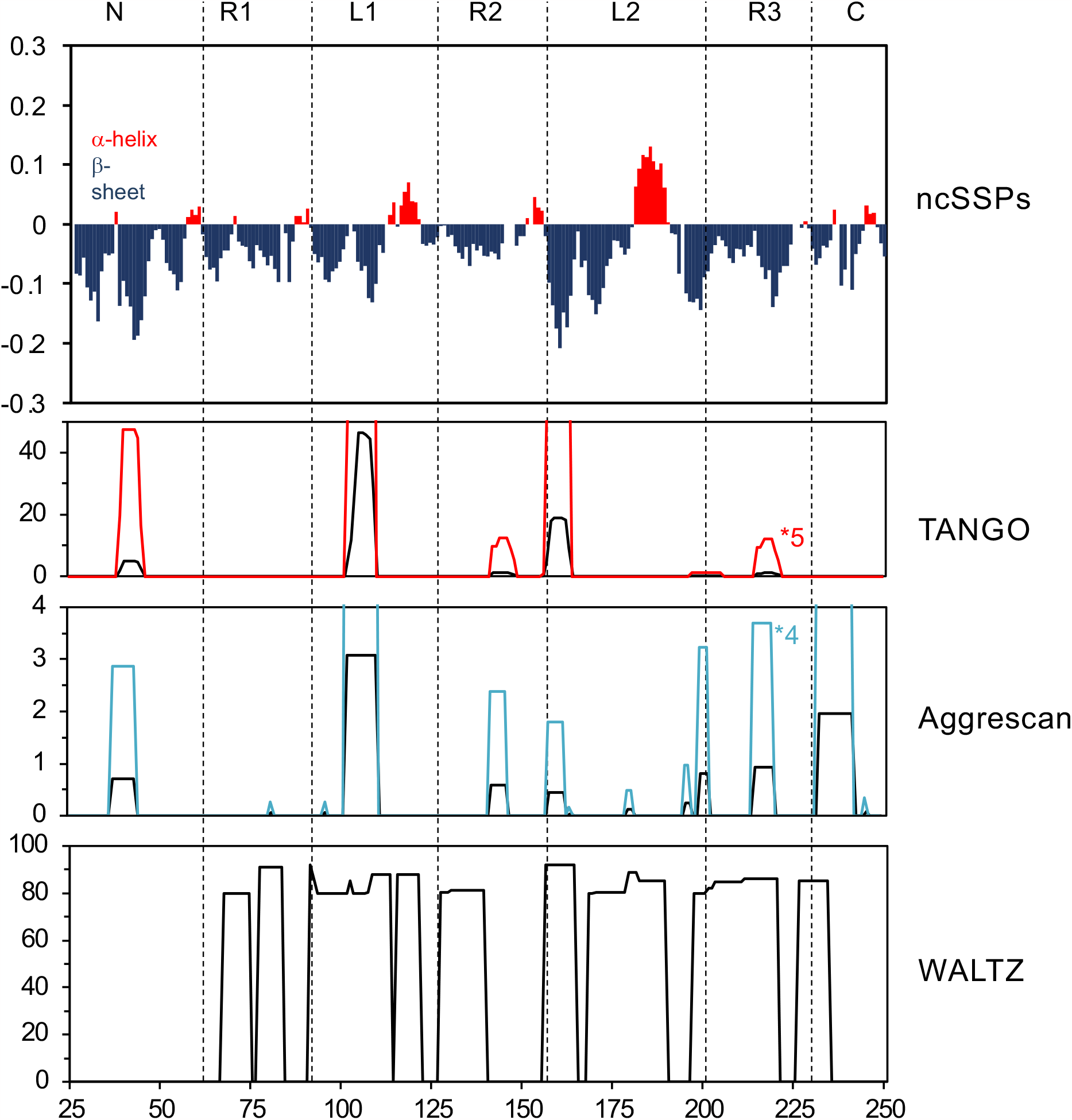

## References

(1) Chiti, F.; Dobson, C. M. Protein Misfolding, Amyloid Formation, and Human Disease: A Summary of Progress Over the Last Decade. In Annual Review of Biochemistry, Vol 86, Kornberg, R. D. Ed.; Annual Review of Biochemistry, Vol. 86; 2017; pp 27–68.

(2) Rubel, M. S.; Fedotov, S. A.; Grizel, A. V.; Sopova, J. V.; Malikova, O. A.; Chernoff, Y. O.; Rubel, A. A. Functional Mammalian Amyloids and Amyloid-Like Proteins. Life-Basel 2020, 10 (9). DOI: 10.3390/life10090156.

(3) Sawaya, M. R.; Hughes, M. P.; Rodriguez, J. A.; Riek, R.; Eisenberg, D. S. The expanding amyloid family: Structure, stability, function, and pathogenesis. Cell 2021, 184 (19), 4857–4873. DOI: 10.1016/j.cell.2021.08.013.

(4) Salinas, N.; Povolotsky, T. L.; Landau, M.; Kolodkin-Gal, I. Emerging Roles of Functional Bacterial Amyloids in Gene Regulation, Toxicity, and Immunomodulation. Microbiology and Molecular Biology Reviews 2021, 85 (1). DOI: 10.1128/mmbr.00062-20.

(5) Levkovich, S. A.; Gazit, E.; Bar-Yosef, D. L. Two Decades of Studying Functional Amyloids in Microorganisms. Trends in Microbiology 2021, 29 (3), 251–265. DOI: 10.1016/j.tim.2020.09.005.

(6) Akbey, U.; Andreasen, M. Functional amyloids from bacterial biofilms - structural properties and interaction partners. Chemical Science 2022, 13 (22), 6457–6477. DOI: 10.1039/d2sc00645f.

(7) Bjarnsholt, T. The role of bacterial biofilms in chronic infections. Apmis 2013, 121, 1–58. DOI: 10.1111/apm.12099.

(8) Chapman, M. R.; Robinson, L. S.; Pinkner, J. S.; Roth, R.; Heuser, J.; Hammar, M.; Normark, S.; Hultgren, S. J. Role of Escherichia coli curli operons in directing amyloid fiber formation. Science 2002, 295 (5556), 851–855. DOI: 10.1126/science.1067484.

(9) Dueholm, M. S.; Petersen, S. V.; Sonderkaer, M.; Larsen, P.; Christiansen, G.; Hein, K. L.; Enghild, J. J.; Nielsen, J. L.; Nielsen, K. L.; Nielsen, P. H.; et al. Functional amyloid in Pseudomonas. Molecular Microbiology 2010, 77 (4), 1009–1020. DOI: 10.1111/j.1365-2958.2010.07269.x.

(10) Romero, D.; Aguilar, C.; Losick, R.; Kolter, R. Amyloid fibers provide structural integrity to Bacillus subtilis biofilms. Proceedings of the National Academy of Sciences of the United States of America 2010, 107 (5), 2230–2234. DOI: 10.1073/pnas.0910560107.

(11) Periasamy, S.; Joo, H. S.; Duong, A. C.; Bach, T. H. L.; Tan, V. Y.; Chatterjee, S. S.; Cheung, G. Y. C.; Otto, M. How Staphylococcus aureus biofilms develop their characteristic structure. Proceedings of the National Academy of Sciences of the United States of America 2012, 109 (4), 1281–1286. DOI: 10.1073/pnas.1115006109.

(12) Khambhati, K.; Patel, J.; Saxena, V.; Parvathy, A.; Jain, N. Gene Regulation of Biofilm-Associated Functional Amyloids. Pathogens 2021, 10 (4). DOI: 10.3390/pathogens10040490.

(13) Turoverov, K. K.; Kuznetsova, I. M.; Uversky, V. N. The protein kingdom extended: Ordered and intrinsically disordered proteins, their folding, supramolecular complex formation, and aggregation. Progress in Biophysics & Molecular Biology 2010, 102 (2-3), 73–84. DOI: 10.1016/j.pbiomolbio.2010.01.003.

(14) Jain, N.; Chapman, M. R. Bacterial functional amyloids: Order from disorder. Biochimica Et Biophysica Acta-Proteins and Proteomics 2019, 1867 (10), 954–960. DOI: 10.1016/j.bbapap.2019.05.010.

(15) Sewell, L.; Stylianou, F.; Xu, Y. Q.; Taylor, J.; Sefer, L.; Matthews, S. NMR insights into the pre-amyloid ensemble and secretion targeting of the curli subunit CsgA. Scientific Reports 2020, 10 (1). DOI: 10.1038/s41598-020-64135-9.

(16) Stelzl, L. S.; Pietrek, L. M.; Holla, A.; Oroz, J.; Sikora, M.; Kofinger, J.; Schuler, B.; Zweckstetter, M.; Hummer, G. Global Structure of the Intrinsically Disordered Protein Tau Emerges from Its Local Structure. Jacs Au 2022, 2 (3), 673–686. DOI: 10.1021/jacsau.1c00536.

(17) Rasmussen, H. Ø.; Kumar, A.; Shin, B.; Stylianou, F.; Sewell, L.; Xu, Y.; Otzen, D. E.; Pedersen, J. S.; Matthews, S. J. FapA is an Intrinsically Disordered Chaperone for Pseudomonas Functional Amyloid FapC. Journal of Molecular Biology 2023, 435 (2), 167878. 10.1016/j.jmb.2022.167878.

(18) Karamanos, T. K.; Kalverda, A. P.; Thompson, G. S.; Radford, S. E. Mechanisms of amyloid formation revealed by solution NMR. Progress in Nuclear Magnetic Resonance Spectroscopy 2015, 88-89, 86–104. DOI: 10.1016/j.pnmrs.2015.05.002.

(19) Gallardo, R.; Ranson, N. A.; Radford, S. E. Amyloid structures: much more than just a cross-beta fold. Current Opinion in Structural Biology 2020, 60, 7–16. DOI: 10.1016/j.sbi.2019.09.001.

(20) Levine, H. THIOFLAVINE-T INTERACTION WITH SYNTHETIC ALZHEIMERS-DISEASE BETA-AMYLOID PEPTIDES - DETECTION OF AMYLOID AGGREGATION IN SOLUTION. Protein Science 1993, 2 (3), 404–410. DOI: 10.1002/pro.5560020312.

(21) Ulamec, S. M.; Brockwell, D. J.; Radford, S. E. Looking Beyond the Core: The Role of Flanking Regions in the Aggregation of Amyloidogenic Peptides and Proteins. Frontiers in Neuroscience 2020, 14. DOI: 10.3389/fnins.2020.611285.

(22) Okamura, E.; Aki, K. Real-time in-situ H-1 NMR of reactions in peptide solution: preaggregation of amyloid-beta fragments prior to fibril formation. Pure and Applied Chemistry 2020, 92 (10), 1575–1583. DOI: 10.1515/pac-2019-1201.

(23) Sewell, L.; Stylianou, F.; Xu, Y.; Taylor, J.; Sefer, L.; Matthews, S. NMR insights into the pre-amyloid ensemble and secretion targeting of the curli subunit CsgA. Sci Rep 2020, 10 (1), 7896. DOI: 10.1038/s41598-020-64135-9.

(24) Santoro, A.; Grimaldi, M.; Buonocore, M.; Stillitano, I.; D’Ursi, A. M. Exploring the Early Stages of the Amyloid A beta(1-42) Peptide Aggregation Process: An NMR Study. Pharmaceuticals 2021, 14 (8). DOI: 10.3390/ph14080732.

(25) Clore, G. M. NMR spectroscopy, excited states and relevance to problems in cell biology - transient pre-nucleation tetramerization of huntingtin and insights into Huntington’s disease. Journal of Cell Science 2022, 135 (12). DOI: 10.1242/jcs.258695.

(26) Fawzi, N. L.; Ying, J. F.; Torchia, D. A.; Clore, G. M. Kinetics of Amyloid beta Monomer-to-Oligomer Exchange by NMR Relaxation. Journal of the American Chemical Society 2010, 132 (29), 9948–9951. DOI: 10.1021/ja1048253.

(27) Fawzi, N. L.; Ying, J. F.; Ghirlando, R.; Torchia, D. A.; Clore, G. M. Atomic-resolution dynamics on the surface of amyloidbeta protofibrils probed by solution NMR. Nature 2011, 480 (7376), 268–U161. DOI: 10.1038/nature10577.

(28) Davies, H. A.; Rigden, D. J.; Phelan, M. M.; Madine, J. Probing Medin Monomer Structure and its Amyloid Nucleation Using 13C-Direct Detection NMR in Combination with Structural Bioinformatics. Scientific Reports 2017, 7 (1), 45224. DOI: 10.1038/srep45224.

(29) Kotler, S. A.; Tugarinov, V.; Schmidt, T.; Ceccon, A.; Libich, D. S.; Ghirlando, R.; Schwieters, C. D.; Clore, G. M. Probing initial transient oligomerization events facilitating Huntingtin fibril nucleation at atomic resolution by relaxation-based NMR. Proceedings of the National Academy of Sciences of the United States of America 2019, 116 (9), 3562–3571. DOI: 10.1073/pnas.1821216116.

(30) Schwarze, B.; Korn, A.; Höfling, C.; Zeitschel, U.; Krueger, M.; Roßner, S.; Huster, D. Peptide backbone modifications of amyloid β (1–40) impact fibrillation behavior and neuronal toxicity. Scientific Reports 2021, 11 (1), 23767. DOI: 10.1038/s41598-021-03091-4.

(31) Chang-Hyeock Byeon, P. C. W., In-Ja L. Byeon, Umit Akbey. Solution-state NMR Assignment and Secondary Structural Propensities of the Full-Length and Minimalistic-Truncated Prefibrillar Monomeric Form of Biofilm-Forming Functional-Amyloid FapC from Pseudomonas aeruginosa. bioRxiv. DOI: 10.1101/2023.01.22.525107.

(32) Norrild, R. K.; Vettore, N.; Coden, A.; Xue, W. F.; Buell, A. K. Thermodynamics of amyloid fibril formation from non-equilibrium experiments of growth and dissociation. Biophysical Chemistry 2021, 271. DOI: 10.1016/j.bpc.2021.106549.

(33) Burmann, B. M.; Gerez, J. A.; Matecko-Burmann, I.; Campioni, S.; Kumari, P.; Ghosh, D.; Mazur, A.; Aspholm, E. E.; Sulskis, D.; Wawrzyniuk, M.; et al. Regulation of alpha-synuclein by chaperones in mammalian cells. Nature 2020, 577 (7788), 127–+. DOI: 10.1038/s41586-019-1808-9.

(34) Wentink, A. S.; Nillegoda, N. B.; Feufel, J.; Ubartaite, G.; Schneider, C. P.; De Los Rios, P.; Hennig, J.; Barducci, A.; Bukau, B. Molecular dissection of amyloid disaggregation by human HSP70. Nature 2020, 587 (7834), 483–+. DOI: 10.1038/s41586-020-2904-6.

(35) Williams, D. M.; Thorn, D. C.; Dobson, C. M.; Meehan, S.; Jackson, S. E.; Woodcock, J. M.; Carver, J. A. The Amyloid Fibril-Forming beta-Sheet Regions of Amyloid beta and alpha-Synuclein Preferentially Interact with the Molecular Chaperone 14-3-3 zeta. Molecules 2021, 26 (20). DOI: 10.3390/molecules26206120.

(36) Piovesan, D.; Walsh, I.; Minervini, G.; Tosatto, S. C. E. FELLS: fast estimator of latent local structure. Bioinformatics 2017, 33 (12), 1889–1891. DOI: 10.1093/bioinformatics/btx085 (acccessed 2/10/2023).

(37) Nagaraj, M.; Ahmed, M.; Lyngso, J.; Vad, B. S.; Boggild, A.; Fillipsen, A.; Pedersen, J. S.; Otzen, D. E.; Akbey, U. Predicted Loop Regions Promote Aggregation: A Study of Amyloidogenic Domains in the Functional Amyloid FapC. Journal of Molecular Biology 2020, 432 (7), 2232–2252. DOI: 10.1016/j.jmb.2020.01.044.

(38) Linding, R.; Schymkowitz, J.; Rousseau, F.; Diella, F.; Serrano, L. A comparative study of the relationship between protein structure and beta-aggregation in globular and intrinsically disordered proteins. Journal of Molecular Biology 2004, 342 (1), 345–353. DOI: 10.1016/j.jmb.2004.06.088.

(39) Conchillo-Sole, O.; de Groot, N. S.; Aviles, F. X.; Vendrell, J.; Daura, X.; Ventura, S. AGGRESCAN: a server for the prediction and evaluation of “hot spots” of aggregation in polypeptides. Bmc Bioinformatics 2007, 8. DOI: 10.1186/1471-2105-8-65.

(40) Louros, N.; Konstantoulea, K.; De Vleeschouwer, M.; Ramakers, M.; Schymkowitz, J.; Rousseau, F. WALTZ-DB 2.0: an updated database containing structural information of experimentally determined amyloid-forming peptides. Nucleic Acids Research 2020, 48 (D1), D389–D393. DOI: 10.1093/nar/gkz758.

(41) Blanch, E. W.; Morozova-Roche, L. A.; Cochran, D. A. E.; Doig, A. J.; Hecht, L.; Barron, L. D. Is polyproline II helix the killer conformation? A Raman optical activity study of the amyloidogenic prefibrillar intermediate of human lysozyme. Journal of Molecular Biology 2000, 301 (2), 553–563. DOI: 10.1006/jmbi.2000.3981.

(42) Dueholm, M. S.; Otzen, D.; Nielsen, P. H. Evolutionary Insight into the Functional Amyloids of the Pseudomonads. Plos One 2013, 8 (10). DOI: 10.1371/journal.pone.0076630.

(43) Byeon, C.-H.; Akbey, Ü. Solution-state NMR Assignment and Secondary Structure Analysis of the Monomeric <em>Pseudomonas</em> Biofilm-Forming Functional Amyloid Accessory Protein FapA. bioRxiv 2023, 2023.2007.2018.549541. DOI: 10.1101/2023.07.18.549541.

(44) Andreasen, M.; Meisl, G.; Taylor, J. D.; Michaels, T. C. T.; Levin, A.; Otzen, D. E.; Chapman, M. R.; Dobson, C. M.; Matthews, S. J.; Knowles, T. P. J. Physical Determinants of Amyloid Assembly in Biofilm Formation. Mbio 2019, 10 (1). DOI: 10.1128/mBio.02279-18.

(45) Bardwell, J. C. A.; Jakob, U. Conditional disorder in chaperone action. Trends in Biochemical Sciences 2012, 37 (12), 517–525. DOI: 10.1016/j.tibs.2012.08.006.

(46) Ikeda, K.; Suzuki, S.; Shigemitsu, Y.; Tenno, T.; Goda, N.; Oshima, A.; Hiroaki, H. Presence of intrinsically disordered proteins can inhibit the nucleation phase of amyloid fibril formation of A beta(1-42) in amino acid sequence independent manner. Scientific Reports 2020, 10 (1). DOI: 10.1038/s41598-020-69129-1.

(47) Ulamec, S. M.; Maya-Martinez, R.; Byrd, E. J.; Dewison, K. M.; Xu, Y.; Willis, L. F.; Sobott, F.; Heath, G. R.; Hawle, P. V.; Buchman, V. L.; et al. Single residue modulators of amyloid formation in the N-terminal P1-region of alpha-synuclein. Nature Communications 2022, 13 (1). DOI: 10.1038/s41467-022-32687-1.

(48) Jumper, J.; Evans, R.; Pritzel, A.; Green, T.; Figurnov, M.; Ronneberger, O.; Tunyasuvunakool, K.; Bates, R.; Zidek, A.; Potapenko, A.; et al. Highly accurate protein structure prediction with AlphaFold. Nature 2021, 596 (7873), 583–+. DOI: 10.1038/s41586-021-03819-2.

(49) Salmon, L.; Nodet, G.; Ozenne, V.; Yin, G. W.; Jensen, M. R.; Zweckstetter, M.; Blackledge, M. NMR Characterization of Long-Range Order in Intrinsically Disordered Proteins. Journal of the American Chemical Society 2010, 132 (24), 8407–8418. DOI: 10.1021/ja101645g.

(50) Taylor, J. D.; Hawthorne, W. J.; Lo, J.; Dear, A.; Jain, N.; Meisl, G.; Andreasen, M.; Fletcher, C.; Koch, M.; Darvill, N.; et al. Electrostatically-guided inhibition of Curli amyloid nucleation by the CsgC-like family of chaperones. Scientific Reports 2016, 6. DOI: 10.1038/srep24656.

(51) Wishart, D. S.; Bigam, C. G.; Holm, A.; Hodges, R. S.; Sykes, B. D. H-1, C-13 AND N-15 Random coil nmr chemical-shifts of the common amino-acids .1. investigations of nearest-neighbor effects. Journal of Biomolecular Nmr 1995, 5 (1), 67–81. DOI: 10.1007/bf00227471.

